# Ant identity determines the fungi richness and composition of a myrmecochorous seed

**DOI:** 10.1101/2023.10.12.562019

**Authors:** Tiago V. Fernandes, Otavio L. Fernandes, Inácio J. M. T. Gomes, Ricardo R. C. Solar, Ricardo I. Campos

**Affiliations:** Programa de Pós-Graduação em Biologia Animal, Universidade Federal dos Vales do Jequitinhonha e Mucuri. Diamantina, Brazil; Programa de Pós-Graduação em Ecologia, Conservação e Manejo da Vida Silvestre. Universidade Federal de Minas Gerais. Instituto de Ciências Biológicas, Belo Horizonte, Brazil; Universidade Federal de Viçosa, Departamento de Biologia Geral, Viçosa-MG, Brazil; Universidade Federal de Viçosa, Departamento de Entomologia, Viçosa-MG, Brazil

**Author notes:** **Author contributions:** All authors have made crucial scientific contributions to the study. Material preparation, data collection and analysis were performed by OLF and TVF. The manuscript was written by TVF. All authors commented on previous versions of the manuscript and approved the final manuscript. All stages were supervised by RIC.

**Keywords:** Seed dispersal, seed handling, leaf-cutting ants, mold, Formicidae, dispersal syndrome

## Abstract

Myrmecochory - seed dispersal by ants - is a mutualistic interaction in which ants attracted by seed appendices take them away from the parental plant location, where seeds usually have better development odds. Not all ant species benefit plants, and the mechanisms of those divergent outcomes are still unclear, especially from the perspective of microbial third parties. Here, we explore the effects of seed manipulation on fungi communities promoted by two ant species with contrasting effects on seed germination and antimicrobial cleaning strategies. We hypothesize that: i) fungi richness is higher in seeds manipulated by *Acromyrmex subterraneus* (species that negatively affect seed germination), followed by unmanipulated seeds and seeds manipulated by *Atta sexdens* (ant species that increase seed germination) and ii) seeds manipulated by *A. sexdens, Ac. subterraneus* and unmanipulated seeds present dissimilar fungi compositions. We identified fungal morphotypes in three groups of seeds: i) manipulated by *A. sexdens*; ii) manipulated by *Ac. subterraneus*; iii) unmanipulated. Seeds manipulated by *Ac. subterraneus* exhibited higher fungal richness than those manipulated by *A. sexdens* and unmanipulated seeds, indicating that the ant species known to impair germination increases the fungal load on seeds. Additionally, we found that *A. sexdens* ants were unable to reduce fungal richness compared to unmanipulated seeds. Furthermore, fungal composition differed among all three treatments. Our results underscore the significance of ant species identity in shaping the fungal communities associated with myrmecochorous seeds. Given the potential influence of microbial infection on seed fate, we suggest considering manipulation strategies when evaluating the overall quality of an ant as a seed disperser.

## Introduction

Mutualisms between plants and insects represent a major topic in ecology and evolution [1]. An important benefit that insects provide to plants is seed dispersal, with myrmecochory (i.e., seed dispersal by ants) standing out as the sole true seed dispersal syndrome among invertebrates, observed in over 11,000 plant species [2–4]. Myrmecochorous seeds are characterized by an oily appendage, called elaiosome, which ants use as a food source [5,6]. While seeds provide food to ants, the latter can increase the reproductive success of plants by manipulating and moving their seeds to places with better development odds [4,7]. Most studies focusing on the advantages of myrmecochory to plants focus on two mechanisms: seed removal distance and deposition site [4]. On the other hand, the effects of ants’ physical and chemical manipulation of seeds have been poorly investigated ([but see 8,9]. Because not all seed disperser ants benefit plants [10,11], comparing ant species is fundamental to understanding the holy scenario on the possible effects of myrmecochory on seed germination, plant development, and consequently, plant population dynamics.

When ants manipulate seeds, they remove the elaiosome or pulp to feed on [8,12]. During this process, ants can also promote seed disinfection by actively removing microorganisms or deploying antimicrobial substances produced mainly by their exocrine glands [9,13]. Ant cleaning strategy prevents the growth of pathogenic fungi on non-myrmecochorous seeds [9,12] and changes the fungi communities surrounding ant nests [14], but ant effect on fungi communities of myrmecochorous seeds has never been tested.

Ant species employ distinct cleaning strategies to safeguard themselves and their colonies against pathogenic microorganisms. [15–17]. Also, the frequency of use and chemical composition of antimicrobial substances secreted by ants glands are highly variable among species [16]. Therefore, each ant species can have different outcomes in their disinfection efficiency against pathogens. It is important to note that ant hygienic strategies also inhibit a variety of microorganisms with no evident harm to ants, such as phytopathogenic fungi. [13,14,18]. Therefore, distinct hygienic strategies among ant species might play divergent roles in the fungi communities present on myrmecochorous seed.

*Atta sexdens* (Linnaeus, 1758) and *Acromyrmex subterraneus* (Forel, 1893) are two sympatric and genetically related species of leaf-cutting ants with distinguishing hygienic strategies [19]. The genus *Atta* immune strategy is based on the production of chemical compounds by exocrine glands, mainly the metapleural and mandibular glands (Fernandez-Marin et al. 2009). On the other hand, ants of the genus *Acromyrmex* have a symbiotic association with *Pseudonocardia*. In this case, the bacteria grow under the ant tegument and produce antibiotics, which act as the primary line of defense against ant pathogenic infections [19–21]. Finally, it is known that the secretion produced by the *Pseudonocardia* bacteria has a smaller spectrum of action against microorganisms when compared to metapleural gland compounds [19].

Those two ant species disperse seeds of the myrmecochorous plant *Mabea fistulifera* Mart.-Euphorbiaceae[22]. *Mabea fistulifera* is a pioneer tree species widely distributed in Brazil and presents a two-phase seed dispersal system (Diplochory). When fruits ripen and open, seeds are ejected onto the ground (up to 8 m, a process known as ballochory), where ants collect and transport them to their nest (Fernandes et al. 2020). Both *A. sexdens* and *Ac. subterraneus* are known to retrieve *M. fistulifera* seeds from the ground and transport them to their nest. In the nest, they consume the elaiosome, and afterwards, they discard the seeds on the soil surface near the nest entrance [7,8,22]. Although those two species have similar dispersal behaviours, they present contrasting outcomes in seed germination [8,19,23]. The manipulation of *M. fistulifera* seeds by *A. sexdens* improves germination rates, while the manipulation of seeds by *Ac. subterraneus* could cause no change or even a decrease in the germination of those seeds [8,23]. Hence, variations in hygienic strategies between *A. sexdens* and *Ac. subterraneus* ants could directly impact microbial communities, including fungi, on myrmecochorous seeds, and potentially affect germination.

Based on this, our main objective was to experimentally compare the fungi richness and composition on seeds manipulated by *Atta sexdens, Acromyrmex subterraneus* and unmanipulated seeds. We tested two non-excluding hypotheses: i) fungi richness is higher in seeds manipulated by *Ac. subterraneus* (species that negatively affect seed germination), followed by unmanipulated seeds and seeds manipulated by *A. sexdens* (ant species that increase seed germination). ii) seeds manipulated by *A. sexdens, Ac. subterraneus* and unmanipulated seeds present dissimilar fungi compositions. By doing so, we provide insight into the role of third-party microbial interactions with ant species in myrmecochory.

## Material and Methods

### Study area and experimental design

We gathered mature yet unripe fruits from ten *Mabea fistulifera* trees in a secondary semi-deciduous Atlantic Forest fragment located in Viçosa, Minas Gerais, Brazil. (20.77°S, 42.96° W). We chose *M. fistulifera* because it has myrmecochorous seeds and is often manipulated by many ant species [22]. Immediately after collecting, we placed the fruits on trays under sunlight for two weeks to promote ripening and seed release. From those fruits, we extracted approximately 1000 seeds. We used those fresh seeds to perform a laboratory experiment developed in the *Laboratório de Formigas Cortadeiras* (LFC), at the Entomology Department of Universidade Federal de Viçosa.

We offered those seeds to two ant species, *Acromyrmex subterraneus* and *Atta sexdens*. Both species are highly attracted by *M. fistulifera* seeds and have well-known seed manipulation behaviours [23]. For the seed manipulation trials, we used four nests of *A. sexdens* and four of *Ac. subterraneus*. Those nests were kept in plastic trays (60 x 40 x 15 cm) with inert powder to prevent ants from escaping and a central chamber (inside a plastic pot) where they cultivated their symbiotic fungus. Each ant nest had been fed only with fresh leaves of *Acalypha wilkesiana* (Euphorbiaceae) and maintained under controlled laboratory conditions (25°C, 12-12 h of light) for two years before we started our experiment. All the nests were kept together in the same room under the conditions described above.

To conduct the experiment, we positioned two Petri dishes (90 x 15 mm) for each experimental nest. One Petri dish was placed inside the nest, while the other was positioned outside the nest, approximately 10 cm away from the nest, ensuring that it remained inaccessible to ants. In each of those Petri dishes, we put 50 *M. fistulifera* seeds. After 48 hours, we collected all seeds that ants had manipulated. We considered as manipulated all seeds that had their elaiosome removed and were discarded by ants in the foraging area outside the Petri dish. From the seeds manipulated by ants, we randomly took a sub-sample of 20 seeds per nest to evaluate the fungi community. For the same purpose, we also took 20 unmanipulated seeds (the ones left outside each experimental nest). Finally, we had three seed treatment groups: i) manipulated by *A. sexdens* (20 seeds per nest = 80 seeds); ii) manipulated by *Ac. Subterraneus* (20 seeds per nest = 80 seeds) and iii) control - unmanipulated seeds left outside of each experimental nest (20 seeds outside of each nest = 160 seeds).

To assess the fungal community on the seeds, we individually placed each seed on sterile Petri dishes (90 x 15 mm) filled with 15 ml of Potato-Dextrose-Agar (PDA) culture medium. Subsequently, we transferred each Petri dish to a Bio-Oxygen-Demand incubator (BOD) set at 25°C, maintaining them for a duration of 28 days. Following this incubation period, we collected samples of the fungi and prepared microscope slides for each distinct fungus morphotype identified within each Petri dish. We identified the fungi to the lowest taxonomic level possible using “The genera of Hyphomycetes” [24] and the website mycobank.org [25]. We employed this method due to its strong specificity in identifying fungi and its widespread use in cost-effective pathogen identification within seeds[26]. Additionally, the Potato-Dextrose-Agar (PDA) medium, being a non-selective growth medium for fungi, renders it suitable for a wide array of fungal species [27].

### Statistical Analyses

To test whether the average richness of fungi differed among the three seed treatment groups, we fitted a generalized linear mixed model (GLMM) with Poisson error distribution using the “lme4” package [28]. We set the seed treatment groups (manipulated by *A. sexdens*, manipulated by *Ac. Subterraneus* and unmanipulated) as the explanatory variable, the number of fungal morphotypes per seed as the response variable, and the experimental nest identity as a random effect. We used the Anova function from the package “mixlm” [29] to check our model’s significance based on the Wald Chi-Squared Test. As we found significance, we assessed differences among manipulation treatments using pairwise contrast analyses [30].

To look for possible changes in the fungi community composition among seed treatment groups, we conducted a Non-Metric Multidimensional Scaling Analysis (NMDS) using “vegan” package [31]. We performed 999 permutations using the number of occurrences of each fungal morphotype per ant nest. To test for possible statistical differences among seed treatment groups formed in NMDS, we use ANOSIM, with 999 permutations using “*labdsv*” package [32]. In both analyses, we adopted the Bray–Curtis dissimilarity index.

We also evaluated if some fungal morphotypes had a stronger association with a given seed treatment group using the Indicator Value Method (IndVal) using the “*indispecies*” package [33]. Indicator values were calculated for each morphotype by comparing the observed distribution of morphotypes with that expected by chance. The significance of the indicator values for each species was evaluated using Multilevel Pattern Analysis with 5000 randomization iterations (significance level = 0.05, [34].

We conducted all analyzes using the R software v 4.1.3 [35] and checked for homoscedasticity and suitability of models using residual analysis. To plot graphs, we used the R packages “*ggplot2*” [36] and “*plyr*” [37].

## Results

We found 24 fungal morphotypes from six families in the three seed treatments (S1 Table). Among those, we could identify 19 morphotypes to the Genera level, as five morphotypes did not have spores for precise identification at any level (*i*.*e*., sterile mycelium). In total, we found 19, 17 and 19 fungal morphotypes on seeds manipulated by *Ac. Subterraneus, A. sexdens* and control, respectively. Despite the similar numbers on the total fungi richness, the average number of fungal morphotypes per seed manipulated by *Ac. Subterraneus* presented around 25% higher species richness (mean ± SE, 2.04 ± 0.12, Chi = 9.1, N = 320, p = 0.02,) than the seeds manipulated by *A*. *sexdens* (1.58 ± 0.08) and control (1.36 ± 0.06), which did not differ from each other (Chi = 2.2034, N = 320, p = 0.13, Fig 1). The non-metric multidimensional scaling analysis (NMDS) showed three distinct fungi community compositions on *M. fistulifera* seeds (manipulated by *Ac. subterraneus*, manipulated by *A. sexdens*, and control, Fig 2). The ANOSIM similarity analysis supported those differences among treatment groups (R^2^ = 0.27, F_2,13_ = 2.408, p = 0.003).

**Fig. 1.**
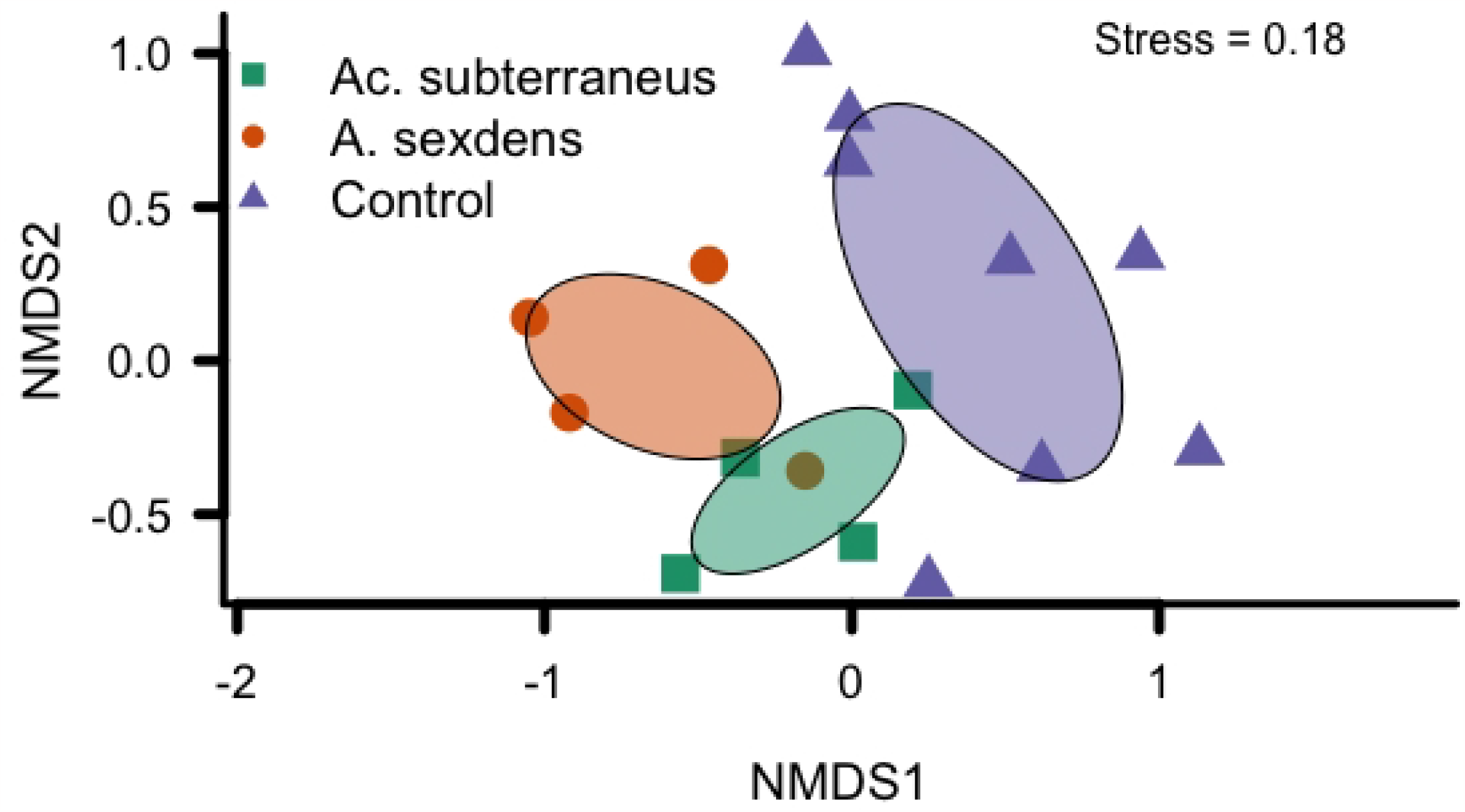
Average richness of fungal morphotypes per seed from treatments: i) seeds previously manipulated by *Acromyrmex subterraneus*, ii) seeds previously manipulated by *Atta sexdens*, iii) seeds not manipulated by ants (Control). Treatments followed by the same letter do not differ statistically among themselves. Points represent the average, bars the standard errors, and shapes the data distribution. *Ac. subterraneus* show a high number of fungal morphotypes than *A. sexdens* and control (Chi = 9.1, N = 320, p = 0.02).

**Fig. 2.**
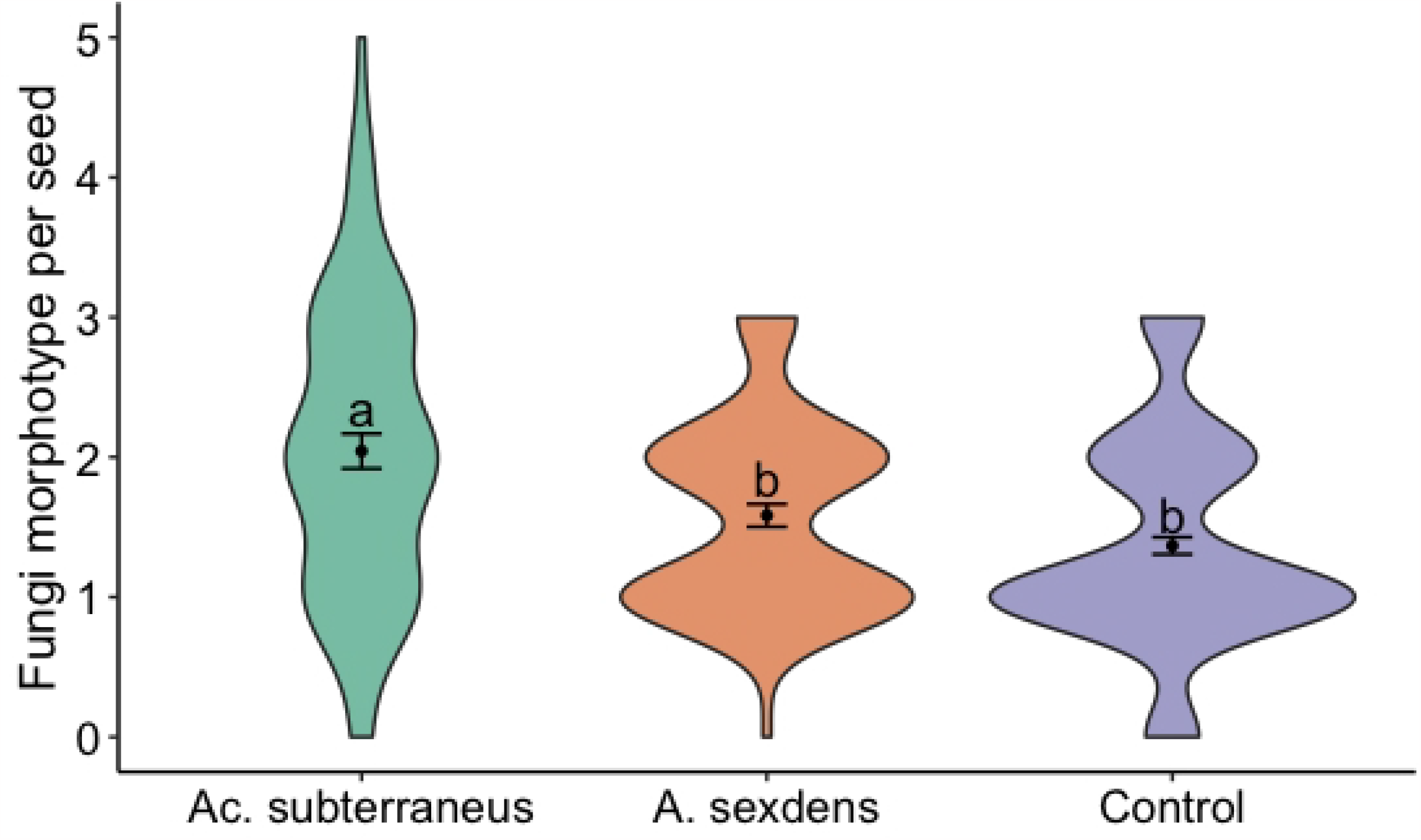
Result of non-metric multidimensional scaling analysis (NMDS): Three groups of fungal morphotypes was formed, one for each treatment i) seeds previously manipulated by *Acromyrmex subterraneus*, ii) seeds previously manipulated by *Atta sexdens*, iii) seeds not manipulated by ants (Control). NMDS was based on the distance from the matrix obtained from the Bray-Curtis index

According to the Species Indicator Value, some fungal morphotypes were more closely associated with a specific seed treatment group. *Penicillium* sp.1 and *Trichoderma* sp.1 were the two most frequent morphotypes on seeds manipulated by *Ac. subterraneus* (IndVal = 0.9, p = 0.005; IndVal = 0.82, p = 0.031, respectively), while the Sterile mycelium sp.2 was mostly associated to the control group (IndVal = 0.92, p = 0.003, S1 Table). Finally, *Fusarium* sp.1 and *Trichothecium roseum* were mainly associated with the two groups of seeds manipulated by ants, regardless of their identity (IndVal = 0.85, p = 0.020; IndVal = 0.79, p = 0.039, respectively). No specific group was associated with *A. sexdens* following IndVal analysis (S1 Table).

## Discussion

Here, we compared seeds manipulated by different ant species and found that *Acromyrmex subterraneus*, a species known to negatively affect the germination of those seeds (Fernandes et al. 2018), exhibited the highest richness of fungi compared to *Atta sexdens* and unmanipulated seeds. We also demonstrate that *A. sexdens* ants were unable to decrease fungal richness when compared to unmanipulated *Mabea fistulifera* seeds. Finally, we showed that fungi composition on these myrmecochorous seeds depends on the ant species which manipulate them. Despite the extensive research on the advantages that ants provide to plants regarding seed removal distance and deposition site, there is relatively limited focus on how ant identity could affect fungi presence on myrmecochorous seeds. Here, our results support the hypothesis that the manipulation of myrmecochorous seeds by different ant species leads to variations in the fungal communities found on seeds. Furthermore, because those two ant species present contrasting effects on seed germination, we suggest that differences in ants’ hygienic strategies contribute to the observed outcomes in seed germination.

We do not support the hypothesis that ants act as seed cleaners by reducing fungal loads on seeds, as suggested for non-myrmecochorous seeds [9,12]. Ants protect themselves and their colonies against infection by releasing antimicrobial substances from external glands or through mutualistic microorganisms [15,19,38]. Mutualist plants may also benefit from this cleaning behaviour by experiencing decreased fungal growth on their seeds [9,12] and reduced pathogenic fungal loads near ant nests where seeds are typically deposited [14,39]. Seeds manipulated by *A. sexdens* showed a similar number of fungal morphotypes compared to unmanipulated seeds, while seeds manipulated by *Ac. subterraneus* had even higher fungal loads. Therefore, leaf-cutting ants do not benefit myrmecochorous seeds by reducing fungal loads on them.

Our results suggest that *Ac. subterraneus* is acting as a seed contaminator instead of a seed cleaner as their manipulated seeds presented more fungal morphotypes than unmanipulated ones. *Acromyrmex subterraneus* is a fungus-growing ant, and its nests have appropriate conditions for developing their symbiotic fungus [40]. However, the same conditions might favour other fungi found growing in the ants’ fungus garden [41]. Furthermore, ants of the genus *Acromyrmex* have their main defences against pathogens based on the antibiotic produced by mutualistic bacteria (*Pseudonocardia* sp.), which has a narrow spectrum of action when compared to glandular secretions commonly used by other ant genera [19–21]. The presence of other fungi inside the ant nest plus the lower spectrum of *Ac. subterraneus* hygienic strategies are likely to explain the increase of fungi infection on seeds manipulated by them compared to the control group. Moreover, *Ac. subterraneus* also changes fungi community on seeds, and *Penicillium* sp.1 and *Trichoderma* sp.1 seem to be favoured by this manipulation. Here, we suggest that the increase in fungi infection and the change in fungi composition make this ant species an inefficient seed cleaner, which might lead to the decrease in the germination observed on *M. fistulifera* seeds manipulated by *Ac. subterraneus* reported by Fernandes *et al* (2020).

Although *A. sexdens* do not actively clean seeds against fungi, their contact during seed dispersal can change fungi composition on the seeds, which could have contributed to the increased germination of *M. fistulifera* after being manipulated by *A. sexdens* seeds [23]. However, it needs to be taken with care as other factors, such as elaiosome removal and seed scarification promoted by ants during the manipulation, could also play a role in seed germination rates [8,42]. Previous studies that have suggested a reduction in fungal infection on seeds through ant manipulation typically assessed only the presence of fungi after germination, without consideration for their specific identity [9,12]. Consequently, the reported fungal infections in prior experiments did not differentiate between fungal attacks that successfully acted as pathogens and those that developed within non-viable seeds. In short, our study indicates that the fungi community present on seeds and the changes in their composition caused by ants are important aspects to consider when measuring myrmecochorous seed fate.

In literature, ant species are classified as high-quality dispersers when they rapidly find and retrieve the seeds to their nests without damaging them [11,43]. For the myrmecochory dispersal syndrome, Giladi (2006) described an ant species as an ideal seed disperser if it retrieves seeds: i) rapidly enough to reduce predation risk, ii) far enough to reduce parental competition, iii) to a suitable site for plant establishment, and iv) without damaging them. Building upon our earlier [7] and the current results, seed manipulation behaviour could be considered an additional factor influencing seed fate because variations in seed manipulation could have consequential effects on seed viability.

Although our results show changes in fungi communities on seeds after being manipulated by different ant species, we have constraints on identifying morphotypes to species. We recognize limitations in our identification of microbial species. Therefore, we suggest further investigation using modern molecular techniques (e.g., next-generation sequencing) to precisely identify fungi, their functional groups and pathogenicity [14,18]. Furthermore, the fine identification of fungi species might be valuable once some fungi lineages could also positively affect seed germination and plant development [44]. Those new methods might give us more detailed information on the perspective of microbial third parties on myrmecochory and their associated ant species.

Our study demonstrates that ants do not decrease fungal loads but fungal composition on myrmecochorous seeds. We further our knowledge about seed dispersal by ants showing that fungi present on seeds depend on the ant species that manipulated them, at least for leaf-cutting ants. In contrast to most studies on myrmecochory, we focus on the differences in ant species considering the cleaning strategies rather than their ability to remove seeds and deposition sites. Our results support the inclusion of one more factor in the myrmecochorous seed fate equation to determine the quality of a seed dispenser, the ant manipulation strategies [9,18,45]. By doing so, we bring new insight into the comprehension of the co-evolutionary paths of the myrmecochory mutualisms, third-part interactions with fungi and their ultimate outcomes on seed fate.

## Acknowledgments

We are grateful to the Departamento de Engenharia Florestal (UFV) for allowing access the area to collect seeds, to Terezinha M. C. Della Lucia’s and Karina Dias Amaral for allowed us to use their ant colonies, and to Meiriele da Silva for help with fungi identification.

## Supporting information

**S1 Table. List of fungal morphotypes found on seeds of *Mabea fistulifera*** after manipulation by *Acromyrmex subterraneus* (80 seeds), *Atta sexdens* (80 seeds), or non-manipulated (control, 160 seeds). “Morphotype Frequency” is the number of seeds contaminated by the fungal morphotypes. The “Indicator value” represents the morphotype association with a given group of seeds.

